# An Expert-Informed Synthetic Animal Data Generator: A Physiology-Consistent Generative Framework for High-Fidelity Animal Digital Twins

**DOI:** 10.64898/2026.04.23.720335

**Authors:** Ali Youssef, Congcong Sun, Tomas Norton

## Abstract

Digital twins are increasingly recognized as a transformative technology for precision livestock farming; however, a major bottleneck in their development remains the scarcity of high-quality, high-granularity physiological data. This study introduces the expert-informed conditional diffusion (EICD) framework, a novel approach to synthesizing high-fidelity metabolic time-series trajectories by embedding mechanistic biological principles directly into the generative process. While traditional generative models often prioritize statistical pattern-matching over biological reality, frequently resulting in physiological hallucinations, the EICD framework utilizes a physiology loss function (PhLF) to act as a form of mechanistic regularization. This guardrail penalizes samples that contradict expert-defined constraints, such as the laws of porcine bioenergetics, effectively steering the model toward a realistic physiological manifold. The framework was validated using an empirical dataset of growing pigs under varying thermal conditions. Quantitative results demonstrate near-perfect statistical distributional fidelity, with the model achieving an average Jensen-Shannon divergence (JSD) of 0.062 and a Kullback-Leibler divergence (KLD) of 0.19. The full EICD model produced a mean energy expenditure (EE) of 284.94 ± 38.70 kJ/kg/day, mirroring the empirical average of 281.33 ± 41.58 kJ/kg/day. In contrast, the standard generative diffusion model (i.e., with no physiology guardrail) exhibited significant distributional drift, yielding a mean EE of 334.41 kJ/kg/day. The biological integrity of the model was further assessed using the biological violation rate (BVR), a novel metric defined as the percentage of generated samples that fall outside the physically possible metabolic boundaries established by species-specific laws. While the standard diffusion model produced frequent biological artifacts, the EICD framework successfully suppressed these hallucinations, ensuring that synthetic trajectories remain strictly grounded in mechanistic laws. Despite these advancements, limitations remain at physiological extremes where individual stochasticity is high. By providing a reliable method for generating physiology-consistent synthetic data, this framework provides a robust foundation for the next generation of animal digital twins.

**Highlights:**

- A novel expert-informed conditional diffusion (EICD) framework is proposed for physiology-consistent synthetic data generation in precision livestock farming.
- A physiology loss function (PhLF) embeds species-specific bioenergetic laws directly into the generative process as a mechanistic guardrail.
- The framework achieves near-perfect distributional fidelity (JSD = 0.062) while suppressing physiological hallucinations (BVR = 0.93%).
- An ablation study confirms that biological consistency is not an emergent property of standard diffusion models but requires explicit mechanistic constraints.
- The framework provides a scalable solution for synthetic data augmentation in precision livestock farming, supporting the 3Rs and enabling high-throughput in silico experimentation.

## Introduction

Precision livestock farming (PLF) relies on the continuous acquisition of high-resolution physiological and behavioural data to enable real-time monitoring, early disease detection , and evidence-based management decisions in commercial animal production systems (Aarts et al., 2024; Berckmans, 2013; Lu et al., 2018; Peña Fernández et al., 2019; Youssef et al., 2013, 2019, 2022). Despite significant advances in wearable sensor technology and automated data collection, the availability of large-scale, high-granularity datasets that capture the full range of an animal’s physiological responses under diverse production conditions remains a critical bottleneck in the development of robust computational models for livestock management. Concurrently, growing societal and regulatory pressure to reduce the burden of invasive animal experimentation has accelerated the adoption of the 3Rs principle (Replacement, Reduction, and Refinement) as a guiding framework for both biomedical and agricultural research (Barré-Sinoussi & Montagutelli, 2015). This shift reflects a broader recognition that computational and data-driven approaches can meaningfully reduce the burden on living animals while advancing biological knowledge. (Tannenbaum & Bennett, 2015).

In this context, the concept of the animal digital twin (ADT) has emerged as a transformative frontier in precision livestock farming and *in silico* experimentation (Youssef et al., 2024). By creating high-fidelity virtual replicas of individual animals, ADTs offer a scalable approach to simulate physiological responses to diverse production stressors, including thermal load, feeding regime, and housing conditions, within a risk-free computational environment, bypassing the need for repeated invasive measurements on live animals. In commercial farming systems, where individual animal monitoring at scale remains prohibitively expensive, ADTs have the potential to transform herd management from reactive to predictive, enabling early identification of welfare deterioration, heat stress events, or sub-clinical disease before observable symptoms emerge. Despite this promise, the realization of a robust ADT in animal production is fundamentally constrained by the availability of massive, high-granularity datasets that can map the continuous trajectories of physiological states across diverse production conditions. The collection of such data through traditional *in vivo* in vivo experimentation is not only cost-prohibitive but also counterproductive to the 3Rs objective of reducing animal use in research. Collecting such data *in vivo* is not only cost-prohibitive but also counterproductive to the goal of reducing animal use.

In the modern computational physiology research, generative machine learning offers a promising path toward resolving this data scarcity issue by producing synthetic physiological trajectories (time-series). Current methodologies for synthetic data generation typically fall into two primary categories: probability distribution techniques and neural network (NN)-based architectures (Ghosheh et al., 2024; Loni et al., 2025).

Probability distribution techniques, such as generative Markov-Bayesian probabilistic modelling (Friston, 2009; Gelfand & Smith, 1990), function by explicitly defining the statistical distributions (e.g., Gaussian or Poisson) that govern the empirical data, leveraging Bayesian inference to incorporate prior domain knowledge. While highly interpretable and capable of hard-coding physical boundaries, these approaches often struggle with high-dimensional, noisy sensor data where the stochasticity of physiological responses cannot be fully captured through rigid mathematical formulas alone.

On the other hand, NN-based techniques, such as generative adversarial networks (GANs), exemplified by frameworks such as AnimalGAN (Chen et al., 2023), diffusion models (DM), and variational autoencoders (VAEs), learn to approximate data distributions through high-capacity architectures (Goodfellow et al., 2020). While these models perform well in capturing intricate temporal patterns of real-world physiological signals, they encounter a fundamental limitation in biological contexts: they prioritize statistical pattern-matching over biological reality. Because they operate as black boxes (Badreldin et al., 2024; Youssef, 2023), they often suffer from “physiological hallucinations,” a term define here as generated data points that violate established mechanistic laws (e.g., energy expenditure falling below the allometric basal metabolic rate) despite appearing statistically plausible, generating data points that violate the fundamental physics/physiology laws such as those of thermodynamics, bioenergetics, and homeostatic regulation. For instance, a synthetic metabolic trajectory might show energy expenditure levels that are physically impossible given the subject’s body mass or ambient temperature.

To bridge the gap between data-driven flexibility and mechanistic thoroughness (Badreldin et al., 2024), there is a critical need for generative frameworks that are not merely data-aware but also “physiology-consistent.” This concept is well recognised in engineering disciplines, where Raissi et al. (Raissi et al., 2019) demonstrated that embedding physical laws as soft constraints into neural network training, or Physics-Informed Neural Networks (PINNs), substantially improves generalization and physical plausibility in fields such as heat and mass transfer. We argue that the same principle is equally critical in biological systems, where mechanistic laws govern feasible physiological states. This paper introduces an expert-informed synthetic data generator that constrains the latent space of generative diffusion models with established physiological and bioenergetic laws. By embedding expert knowledge into the architecture, we ensure that the synthetic outputs serve as a robust, high-fidelity foundation for the next generation of animal digital twins, moving beyond mere statistical accuracy toward biological grounding.

The remainder of this paper is organized as follows: Section 2 (Methodology) presents the EICD framework formulation, including the conditional diffusion model, the physiology loss function, and the expert-defined conditioning inputs. Section 3 (Experiment) describes the empirical porcine dataset, the *in silico* experimental design, and the evaluation metrics. Section 4 (Results and Discussion) presents the statistical, temporal, and physiological fidelity results together with the ablation study findings and their implications for animal digital twin development. Finally, Section 5 (Conclusions and Future Work) summarizes the key contributions and outlines directions for future research.

## Methodology

In this section, we introduce the generic formulation of the expert-informed conditional diffusion (EICD) framework for generating physiology-consistent synthetic data. To demonstrate the framework’s utility, we focus on the generation of metabolic energy expenditure (*EE*) trajectories in pigs across different ages. The measurement of *EE* traditionally requires sophisticated laboratory equipment, such as indirect calorimetry respiration chambers, and extensive animal preparation, making it a prime candidate for high-fidelity synthetic data augmentation.

### Expert-Informed Conditional Diffusion (EICD) Framework

The core idea of this work is the development of the expert-informed conditional diffusion (EICD) framework. This architecture is inspired by the paradigm of physics-informed machine learning (PIML), which integrates seamlessly data and domain-specific mechanistic models into deep learning architectures to ensure that model outputs remain physically consistent (Karniadakis et al., 2021). In the presented approach, we adapt this philosophy to the biological domain by constraining a conditional diffusion model (Higham et al., 2025; Panagiotakopoulos et al., 2025) with the foundational laws of physiology and bioenergetics.

The EICD framework utilizes a diffusion-based generative architecture. The selection of diffusion model (DM) over traditional GANs (e.g., TimeGAN (Brophy et al., 2023)) or VAEs is based on established literature in time-series synthesis (Yi et al., 2024). Diffusion models offer superior training stability and are inherently resistant to mode collapse (Ho et al., 2020; Yi et al., 2024), a common failure in GANs when modelling multi-modal biological signals. By iteratively reversing Gaussian noise through a conditioned denoising process, the model preserves the high-dimensional manifold topography of metabolic trajectories more effectively than the latent-space averaging characteristic of VAEs (Yi et al., 2024).

#### The Conditional Diffusion Model

The EICD frameworks is formulated as a conditional generative process that learn to map latent noise distribution into a structured physiological manifold. Basically, the framework is built upon the principle of denoising diffusion probabilistic models, which define a generative process as the reverse Markov chain that gradually destroys data by adding Gaussian noise (Ho et al., 2020). However, the main innovation here lies in the transition from a purely data-driven model to a physiology-constrained conditional architecture. The EICD framework treats the synthesis of physiological signals as a conditional denoising process (Panagiotakopoulos et al., 2025) over a temporal sequence. Let **x**_0_ = {*x*_0,1_, *x*_0,2_, … , *x*_0,*N*_} be an empirical time-series of length *N*. The EICD framework generates this trajectory based on an empirical conditioning matrix **c** ∈ ℝ^*N*×*D*^ of exogenous signals (e.g., ambient temperature, physical activity, and body mass).

##### a. The forward diffusion process (the noising)

The forward process *q*(·) gradually introduces Gaussian noise into the original empirical time-series signal **x**_0_ ∈ ℝ^*N*×1^, the *EE* in our case here, over T discrete diffusion timesteps. This follows a fixed Markov chain (Panagiotakopoulos et al., 2025) where each step is defined by a noise variance schedule *β*_*t*_, where *β*_*t*_ ∈ (0,1):

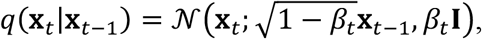

where the **x**_*t*_ is latent noisy times series at step *t*, **x**_*t*−1_ is the previous latent state, 𝒩 denotes a Gaussian (normal) probability distribution, and *β*_*t*_**I** is the covariance matrix (Higham et al., 2025).

As *t* → *T*, the original temporal structure and the underlaying physiological manifold of the original empirical signal **x**_0_ is corrupted, resulting in a sequence of isotropic (direction-independent) Gaussian noise **x**_T_. Using the reparameterization trick, we can sample **x**_*t*_ at any arbitrary timestep *t* directly:

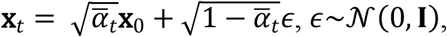

where 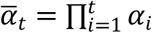 and *α*_*i*_ = 1 − *β*_*i*_ (for more detailed mathematical formulations see (Higham et al., 2025; Yi et al., 2024)).

##### b. The reverse diffusion process (the denoising)

The reverse process *p*_*θ*_(**x**_*t*_|**x**_*t*−1_, **c**), which is basically a denoising backbone of the EICD framework. The objective here is to iteratively remove noise from the latent state to reveal the underlying physiological manifold. To handle the intricate temporal dependencies of the physiological time-series the reverse process utilizes a Gated Recurrent Unit (GRU) as a backbone architecture. The denoising backbone comprises three components: a time-step embedding multi-layer perceptron (MLP; Linear (1→128) → Gaussian error linear unit (GELU) → Linear (128→128)); a single-layer conditioning gated recurrent unit (GRU; input = 3, hidden = 128) that encodes the conditioning context **c**; and a two-layer denoising GRU (input = 257, hidden = 128) that iteratively refines the noisy EE signal. A final linear layer (128→1) maps the GRU output to the predicted noise component.

The reverse process starts from a pure Gaussian noise vector **x**_T_, resulted from the forward diffusion step, and move backwards through T steps. At each step *t*, the GRU-based network *ϵ*_*θ*_(**x**_*t*_, *t*, **c**) predicts the noise component *ϵ* within the current signal **x**_*t*_.

This iterative refinement allows the model to go from a state of total entropy, where all physiological structure has been destroyed and the signal is indistinguishable from pure Gaussian noise, (**x**_*T*_) to a highly structured physiological trajectory (time-series).

In standard conditional diffusion models (Zhou et al., 2024), the following diffusion loss function is defined:

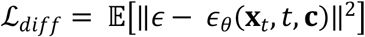

The objective of the diffusion loss function (ℒ_*diff*_) is to ensure that the model learns the statistical distribution of the data, but it does not guarantee that the generated physiological trajectories respect physiological boundaries. Importantly, this objective imposes no explicit constraints on the parameter space, making it agnostic to physiological feasibility.

#### The integration of the expert knowledge

The primary innovation of the EICD framework lies on moving beyond purely statistical learning by embedding expert’s knowledge through two specific channels: a) the conditioning inputs definition and the b) the physiological laws definition:

##### a. Expert-defined conditioning inputs

The proposed framework utilizes an expert-defined conditioning matrix **c** ∈ ℝ^*N*×*D*^ that represents the driving forces and predictors for the target physiological signal (**x**_0_). In this specific case, the target signal is the metabolic energy expenditure (*EE*) and **c** is a set of mechanistic drivers:

- Physical activity, represented by the overall dynamic body acceleration (*ODBA*), which provides the kinetic context for energy demand, serving as a robust proxy for movement-induced thermogenesis (Schrama et al., 1996; Youssef et al., 2020).
- Ambient temperature (*T*_*a*_), which defines the thermoregulatory burden by identifying deviations from the subject’s thermoneutral zone (Bartali et al., 1999; Speakman, 1999).
- Body weight (*BW*), which anchors the allometric baseline of the subject, ensuring the energy requirements are proportional to the animal’s mass and age (Leiva & Schramski, 2022).

##### b. Physiological laws integration (the mechanistic anchor)

To ensure physiological consistency of the generated time-series, we introduced an additional loss function to the EICD framework or the physiology loss function (PhLF) which is mathematically structured to enforce the following bioenergetic laws:

- The lower bound (*L*), represents the minimum possible basal metabolic requirements, ensures the generated synthetic data does not violate the Kleiber’s law (or other laws defined by the expert), which dictates that the basal metabolic rate scales allometrically with body weight (Leiva & Schramski, 2022; National Research Council (NRC), 2012):

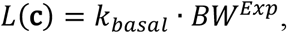

where *k*_*basal*_ is the species-specific metabolic constant and *Exp* is the allometric exponent. This term prevents the model from generating impossible energy states during low activity levels.
- The upper bound (*U*), which represents the maximum physiological energy expenditure achievable under specific environmental and physical conditions. In this work, we have defined *U* to follow the principle of energy balance equations as (note: this formulation is intentionally simplified for the growing-pig proof-of-concept context; anabolic energy costs such as muscle growth or tissue deposition, reproductive/production energy partitioning, and interaction terms between stressors are not explicitly modelled here, but the framework is designed to be extensible by the expert to include such terms as warranted by the target application):

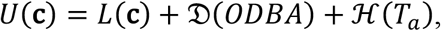

where the term *f*(*ODBA*), activity component (𝒟: the activity energy cost function), represents the energy cost of movement, which is modelled as a function of *ODBA* and *BW* as:

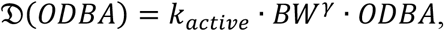

where *k*_*active*_ is the activity coefficient and *γ* is a scaling factor for the energy cost of movement, which is often scales closer to isometric (*BW*^1.0^), i.e., it is directly proportional to the amount of muscle mass (Halsey, 2016; Schrama et al., 1996).

The term ℋ(*T*_*a*_) accounts for metabolic heat production required to maintain homeothermy when *T*_*a*_ deviates from the thermoneutral zone. This relationship can be modelled using the classical Scholander-Irving curve, which defines a U-shaped metabolic floor where energy expenditure is minimal within a specific temperature range and increases linearly as the subject experiences cold or heat stress (Scholander et al., 1950). This relationship us often expressed as a piecewise linear function. However, such function is, kink, non-differentiable at the critical temperature thresholds, the lower critical *T*_*lc*_ and upper critical *T*_*uc*_ temperatures. This can limit gradient-based optimisation in neural network. Therefore, in this work, we utilize the “softplus” function (Zheng et al., 2015), defined as: 𝒮_*plus*_(*x*) = ln(1 + *e*^*x*^), then:

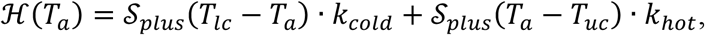

where *k*_*cold*_ and *k*_*hot*_ are thermogenic coefficients represent the rate of increase in metabolic energy expenditure for every degree the ambient temperature deviates out of the animal’s comfort zone.

To enforce biological consistency, the PhLF quantifies deviations from the dynamic validity boundary defined by *L*(**c**) and *U*(**c**). Then a rectified linear unit (ReLU) is employed as a penalty operator. The PhLF here acts as a ‘physiological guardrail’ that only penalize the denoising model when the reconstructed signal 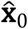 deviates outside the permissible physiological range. The objective of the PhLF is formulated as the expectation of the weighted sum of violations across the sequence *N*:

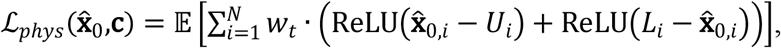

where *w*_*t*_ = (1 − *t*⁄T) is a time-dependent weighting factor that prioritizes physiological consistency during the initial denoising steps, ensuring pruning unrealistic trajectories before the final statistical refinement. The rationale for this time-dependency is that early denoising steps (high noise level, high t) are where the model is most prone to generating trajectories that diverge from the physiological manifold; by applying stronger guardrail penalties at these steps and progressively relaxing them as t → 0, the framework ensures coarse physiological plausibility is established first, after which fine-grained statistical refinement proceeds unimpeded. The clean signal 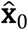 is estimated at each step *t* (Ho et al., 2020) as follows:

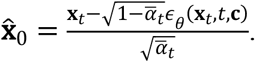

The total EICD loss is a hybrid objective (Karniadakis et al., 2021; Listou Ellefsen et al., 2023; Wang et al., 2025):

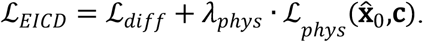

where the term *λ*_*phys*_ is regularisation weight hyperparameter that controls the trad-off between the two competing objectives. A smaller *λ*_*phys*_ prioritises statistical fidelity, allowing the model to capture the underlaying patterns of the empirical data, whereas a larger *λ*_*phys*_ forces the model to strictly follows the expert’s suggested physiological laws.

The EICD framework enables field experts to enforce biological realism by directing the model during the training phase through a physiological guardrail. As listed in Table 1, the expert can constrain the framework using two distinct sets of parameters:

**Table 1.**
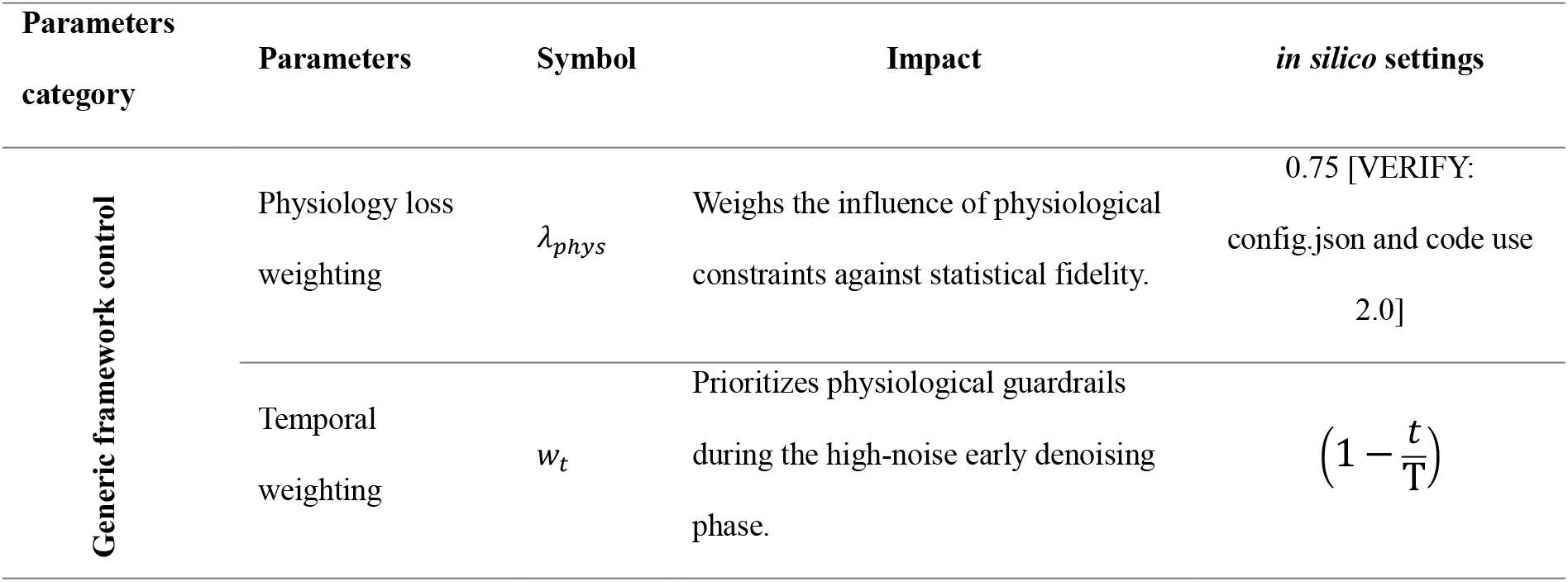
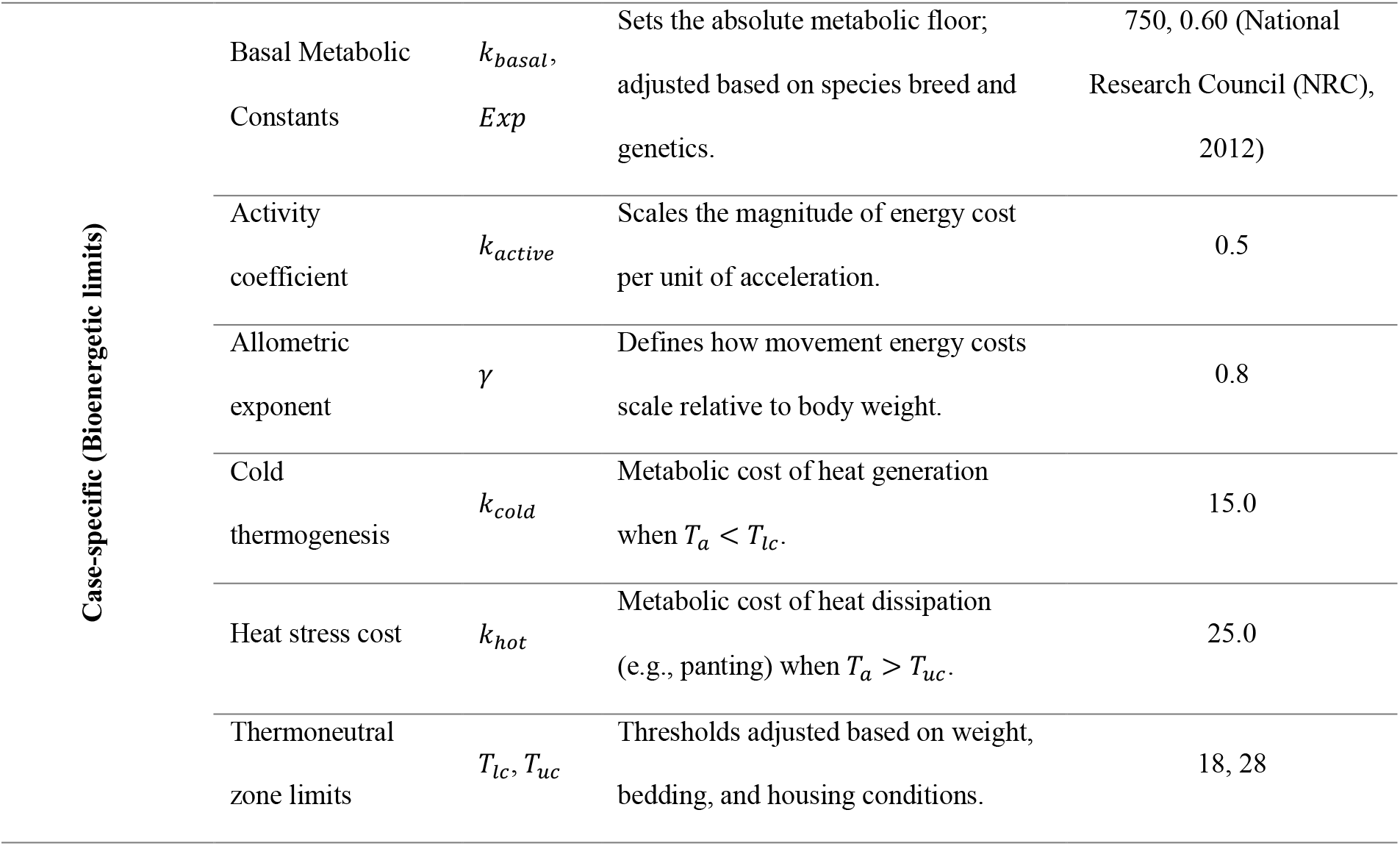
Summary of expert-informed parametric configuration of the EICD architecture. The configuration includes generic hyperparameters for the framework control alongside case-specific bioenergetic parameters driven from porcine metabolic laws. The in silico settings represent the specific parametric values employed during the high throughput in silico experiments.

- Framework control parameters: these EICD generic hyperparameters (e.g., *λ*_*phys*_ and *w*_*t*_) allow the expert to govern the trade-off between following the empirical data patterns or the fundamental biological laws.
- Case-specific physiological limits: these bioenergetic constants (e.g., allometric, activity, and thermoregulation parameters) define the physiological boundaries specific to the animal species and the in silico experimental conditions.

By tuning these parameters, the expert ensures that the generative process is restricted to a permissible validity range, resulting in synthetic data that is both statistically realistic and physiologically grounded.

### Experiment

To evaluate the performance of the EICD framework on empirical data, a controlled longitudinal study was conducted involving porcine subjects. This setup provided the necessary high-fidelity metabolic and behavioural benchmark to test the framework’s ability to generate biologically valid synthetic time-series under varying environmental and physiological conditions. Energy Expenditure (EE) was selected as the primary metric for validation because it serves as a fundamental, dynamic, and real-time indicator of an animal’s physiological state and its interaction with the surrounding environment (Aarts et al., 2024). The focus on EE is further justified by the critical limitations inherent in current gold standard measurement techniques, which hinder the development of robust animal digital twins. Conventional indirect calorimetry (IC) systems, though accurate, are too expensive and require restrictive metabolic chambers that disrupt natural movement and social behaviours. Other techniques, such as doubly labeled water (DLW), provides a total energy budget but lacks the temporal resolution required to capture dynamic fluctuations or to correlate metabolic costs with specific activities (Butler et al., 2004; Heetkamp et al., 2015). These constraints create a significant challenge to continuous, real-time monitoring in real-world settings. By utilizing EE for validation, the EICD framework demonstrates its capacity to fill this technological gap, offering a non-invasive and economically viable.

#### Animal experiment and dataset

The study utilized eight recently weaned female piglets (TN70 x Tempo breed), obtained at four weeks of age. The animals were housed at the CARUS, the animal core facility of Wageningen University & Research, in group pens featuring concrete floors, straw bedding, and environmental enrichment. The ambient temperature was maintained at a constant 22 °C, and pigs were provided ad libitum access to a commercial diet and water. Prior to the formal experiment, pigs underwent a habituation period 6 weeks, to acclimate and train to wearing sensors and isolation within a Climate Respiration Chamber (CRC). All animal handling and experimental procedures followed ethical research standards and received approval from the IVD of Wageningen University, with project number 2022.W-0031.001.

The dataset was generated through three experimental rounds conducted along 10 weeks period to capture metabolic data across different growth stages as shown in Table 2.

**Table 2.** Overview of experimental rounds, pig age, and average metabolic weight across the study duration.

| Experimental round | Pig age (weeks) | Avg. metabolic weight (kg) |
| --- | --- | --- |
| Round 1 | 16-17 | 30.38 ($\pm 1.30$) |
| Round 2 | 21-22 | 38.66 ($\pm 3.10$) |
| Round 3 | 25-26 | 46.14 ($\pm 1.64$) |

During each experimental round, pigs were kept individually in the CRC for three-hour sessions where the ambient temperature (*T*_*a*_) was systematically manipulated to observe thermoregulatory responses. The protocol included both a step-down (cold stress) and step-up (heat stress) phase:

a. Baseline: The CRC was initially set at a constant 22 °C for one hour.
b. Transition: The temperature was adjusted to either 32 °C (heat stress) or 12 °C (cold stress) over a 0.5-hour period.
c. Thermal challenge: The temperature was maintained at the target (32 °C or 12 °C) for one hour.
d. Recovery: the changer returned to 22 °C throughout a final 0.5-hour period.

To ensure the dataset contained enough animal movement data for the activity-based parameters, pigs were kept active through environmental enrichment. Subjects had unrestricted access to water and were provided with paper bags and tube-shaped toys that dispensed snacks, such as peanuts, corn, and dried fruits, to stimulate physical activity throughout their stay in the CRC. To determine metabolic heat expenditure and quantify physical activity, the CRC continuously monitored environmental and physiological variables, including ambient temperature, relative humidity, and gas exchange data (CO_2_). Simultaneously, physical activity was captured via integrated sensors measuring three-axis acceleration at 1 Hz. To isolate dynamic movement from static gravity, a 32-sample sliding window rolling mean was applied to each axis to calculate the *ODBA*.

The final dataset, used for the training of the EICD, comprises 25,718 synchronized observations captured across the longitudinal experimental rounds. The raw sensors data collected at different frequencies were synchronized at a 1-minute sampling rate.

Global statistics of the used longitudinal dataset are summarized in Table 3.

**Table 3.** Statistical summary of the empirical data used for EICD framework validation, showing the distribution of environmental, physical, and metabolic variables for the used porcine subjects (N = 8) across three experimental rounds.

| Parameter | Mean | Std. Dev. | Min | Max |
| --- | --- | --- | --- | --- |
| ***EE* (kJ/kg/day)** | 281.33 | 41.58 | 176.97 | 382.36 |
| ***OBDA* (g)** | 5.26 | 3.24 | 0.02 | 23.49 |
| ***T<sub>a</sub>* (°C)** | 14.66 | 3.92 | 11.89 | 32.79 |
| **BW (kg)** | 38.37 | 6.35 | 30.00 | 46.00 |

#### *In silico* experiment and evaluation

After the EICD framework was trained on the longitudinal porcine dataset, it was deployed in a controlled simulation environment to generate and evaluate synthetic metabolic data. It should be noted that all eight animals were used for model training, and no held-out animal data was reserved for testing; the evaluation presented here is therefore conducted purely through in silico generation, assessing the statistical and physiological fidelity of synthetic trajectories against the full empirical distribution rather than against individual held-out subjects. This approach was chosen to maximize the training signal given the limited dataset size, and its implications for out-of-sample generalization are discussed in the limitations section. This includes generating synthetic *EE* trajectories conditioned on the predefined conditioning variables.

The *in silico* experiment employed a high-throughput generation protocol designed to capture the full variance of the porcine metabolic manifold. The protocol consisted of *R* = 100 independent simulation runs, with each run generating *M* = 1000 *EE* trajectories.

Given a sequence length of *K* (representing a 30-minute window at 1-minute resolution), the experiment resulted in a total synthetic population (𝒟_*syn*_) of 3 × 10^6^ data points:

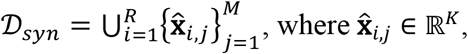

where each individual trajectory 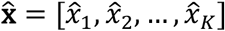 is generated via the stochastic reverse diffusion process. This process is conditioned on the expert-informed conditioning matrix **c** ∈ ℝ^*K*×3^. The generation follows the conditional probability distribution learned by the model: 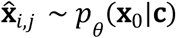.

The *in silico* parametric settings for the expert-informed inputs used in this work are given in Table 1.

In this work, all computational experiments were conducted on the Google Colab platform, utilizing a high-RAM environment equipped with an NVIDIA A100-SXM4-40GB GPU. This hardware configuration provided the 40 GB of HBM2 memory necessary for efficient parallelization of the diffusion reverse steps and the processing of high-frequency multi-sensor pig data. The software architecture was developed in Python 3.10+, primarily leveraging the PyTorch library for the construction and optimization of the deep learning architecture. The diffusion modelling framework utilized a denoising diffusion probabilistic model (DDPM) objective, where training was optimized using the Adam optimizer with a learning rate of 2 × 10^−4^ and a linear noise schedule starting from *β*_*start*_ = 10^− 4^ to *β*_*end*_ = 0.02. Training ran for 500 epochs with a batch size of 128, gradient clipping at max-norm 1.0, and GELU activation in the time-step embedding MLP.

##### a. Ablation study

Rather than benchmarking against external baselines, the primary scientific question of this work, which is isolating the contribution of the physiology guardrail and conditioning matrix to biological plausibility, is best addressed through systematic component-level deconstruction (ablation study), since no existing published method integrates the same combination of physics-informed constraints and conditional diffusion-based generation for physiological time-series in animal systems. Through this systematic deconstruction of the EICD it is possible to isolate the impact of the physiology guardrail (ℒ_*phys*_) and the conditioning matric (**c**) on the model’s ability to generate physiologically viable data. The three variants (ℳ_*full*_, ℳ_*no_phys*_, and ℳ_*uncond*_) isolate each component’s contribution in a controlled manner that external comparisons could not provide, since no existing published method integrates the same combination of physics-informed conditioning and diffusion-based generation for porcine metabolic trajectories.

The full EICD framework (ℳ_*full*_) was benchmarked against two ablated variants as follows:

- ℳ_*full*_: the complete EICD framework, where the *λ*_*phys*_ > 0. This model integrates both expert defined conditioning matrix (**c**) and the physiology guardrail.
- ℳ_*no_phys*_: the physiologically unconstrained diffusion model, where the physiology guardrail is deactivated by setting *λ*_*phys*_ = 0. This model is trained purely on the statistical reconstruction of the empirical data (ℒ_*diff*_).
- ℳ_*uncond*_: the unconditioned diffusion model, where the conditioning matrix (**c**) is nullified during training.

#### b. Evaluation metrics

The quality of the synthetic data generated by the full EICD and each of the ablated variant was evaluated through a combination of point-to-point accuracy, distributional similarity, and physiological consistency. Statistical accuracy was quantified using mean absolute error (MAE) to quantify the deviation between synthetic *EE* and real-world measurements. The distributional similarity was evaluated via two metrics, the Kullback–Leibler (KL) divergence, which measures the information loss when approximating the empirical metabolic distribution, and the Jensen-Shannon divergence (JSD), which provides a symmetrical, normalized similarity score between 0 and 1. To qualitatively validate these results, *t*-distributed stochastic neighbour embedding (*t*-SNE) was employed to visualize the high-dimensional overlap between real and generated data manifolds (Maaten & Hinton, 2008). The temporal fidelity of the generated trajectories was assessed using two additional metrics, the dynamic time warping (DTW), which measures shape similarity by finding the optimal alignment between real and synthetic sequences independently of minor timing offsets (Keogh & Ratanamahatana, 2005; van Veen et al., 2025), providing a more flexible measure of waveform fidelity than point-to-point error (MSE) alone, and normalised cross-correlation, which quantifies the degree of temporal synchrony between the real and synthetic signals and identifies any systematic temporal lag in the model’s metabolic response. Furthermore, a physiological consistency check was performed by calculating the boundary violation rate (BVR), which is the percentage of synthetic data points that falling outside the expert-defined physiological guardrail, ensuring the generated values did not violate the physiological metabolic floors or ceilings. The BVR is given by:

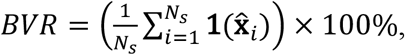

where *N*_*s*_ is the number of synthetic data points and **1**(·) is an indicator function identifies sequences containing any violation of the boundaries, *L*(**c**) and *U*(**c**), of the physiological guardrail:

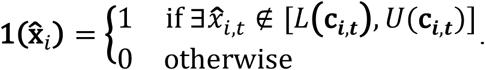

## Results and Discussions

The following section presents a comprehensive evaluation of the EICD framework, analysing its capacity to generate high-fidelity synthetic metabolic trajectories. The evaluation follows a multilevel approach, first, we present the statistical fidelity of the full EICD framework; second, we demonstrate its physiological integrity compared to empirical benchmarks; third, we provide insights from the ablation study. Finally, we discuss the broader implications for the animal digital twin development.

### Overall evaluation of the EICD statistical fidelity

The primary objective of the EICD framework is to capture the underlying probability distribution of the porcine energy expenditure (*EE*) while maintaining high structural similarity to empirical observations. As shown in Table 4, the ℳ_*full*_ model achieved a mean EE of 284.94 ±38.70 kJ/kg/day, showing a negligible deviation from the real data average of 281.33 ±41.58 kJ/kg/day

**Table 4.**
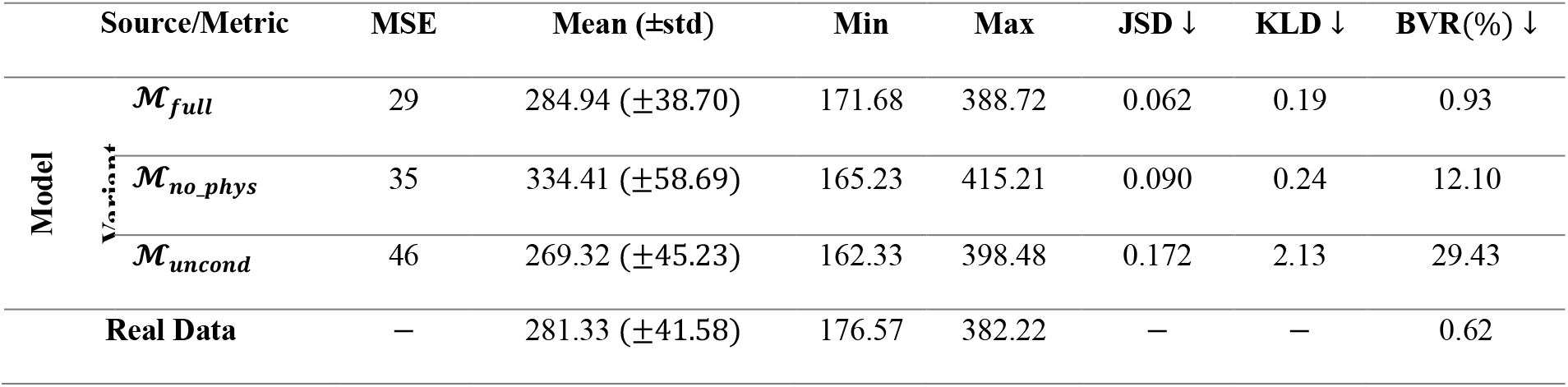
Comparative analysis of generative performance and physiological integrity across model variants. Values represent the mean statistical and biological metrics calculated over R = 100 independent simulation runs (M = 1,000 trajectories per run) against the ground-truth empirical dataset.

The statistical fidelity is further demonstrated by the distributional overlap between real and synthetic data visualized in Figure 1A. The synthetic data density closely mirrors the multi-modal nature of the empirical data, capturing both the basal metabolic peaks and the extended tails associated with high activity thermogenesis. This is also complemented by the violin plot in Figure 1B, which illustrates that the interquartile range and the overall spread of the synthetic population are virtually identical to the real population.

**Figure 1.**
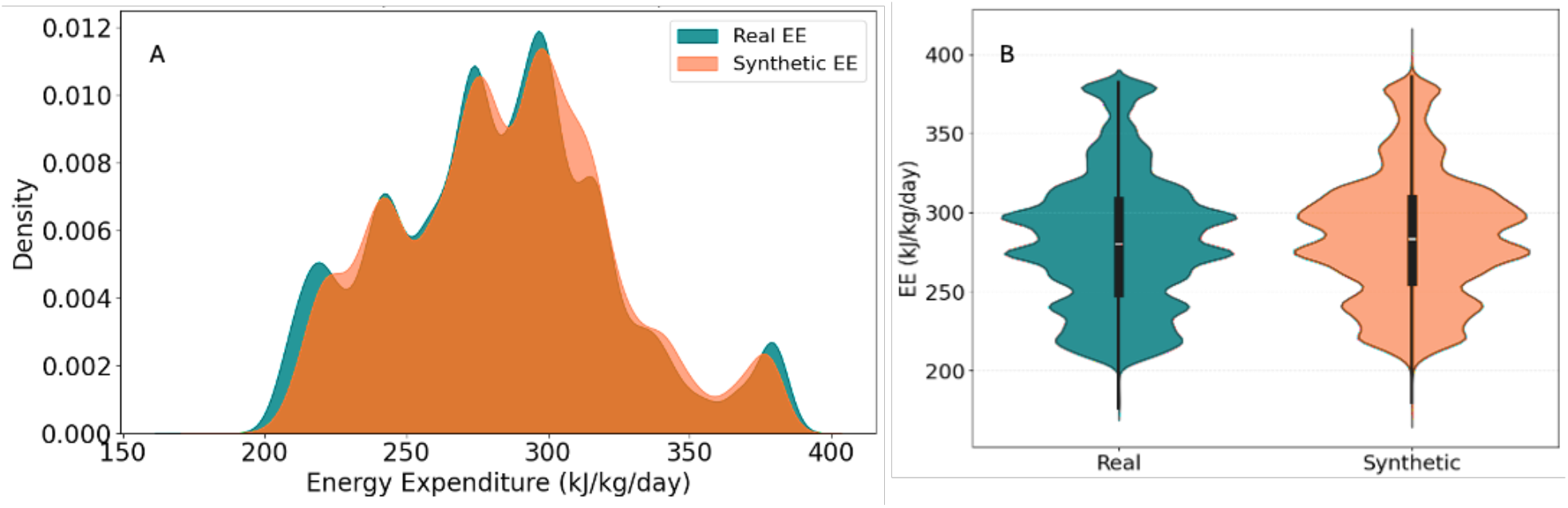
Comparative analysis of population-level Energy Expenditure (EE) fidelity. (A) Kernel density estimate (KDE) demonstrating the overlap of metabolic probability signatures between real and synthetic populations. (B) Violin plots displaying the alignment of medians and interquartile ranges.

This statistical fidelity is quantitatively supported by an average JSD of 0.062 and KLD of 0.19 for the full EICD model (ℳ_*full*_) variant (Table 4). These values indicate that the physiological guardrail does not restrict the model’s generative diversity but rather steer it towards a realistic physiological manifold. In contrast, the ℳ_*no_phys*_ variant showed significant distributional drift, with a mean *EE* of 334.41 kJ/kg/day. This systematic overestimation indicates that a purely statistical diffusion model lacks the mechanistic regularization required to penalize energy values that diverge from the species-specific metabolic baseline.

To evaluate whether the generated 30-minute trajectories (*K* =30) preserve the high-dimensional structural relationships of real metabolic sequences, we employed *t*-SNE visualization. As illustrated in Figure 2, the synthetic trajectories exhibit an overlap with the empirical manifold. The absence of isolated synthetic clusters suggests that the EICD framework avoids mode collapse and does not generate outlier behaviours that fall outside the known porcine metabolic repertoire. To quantify the degree of manifold overlap observed in Figure 2, a coverage score was computed using a k-nearest neighbour (*k*=5) framework in the *t*-SNE embedding space. The coverage score, defined as the percentage of real trajectory points that have at least one synthetic neighbour within their local neighbourhood radius, was 87%, indicating that the EICD framework successfully reconstructs most of the empirical metabolic manifold.

**Figure 2.**
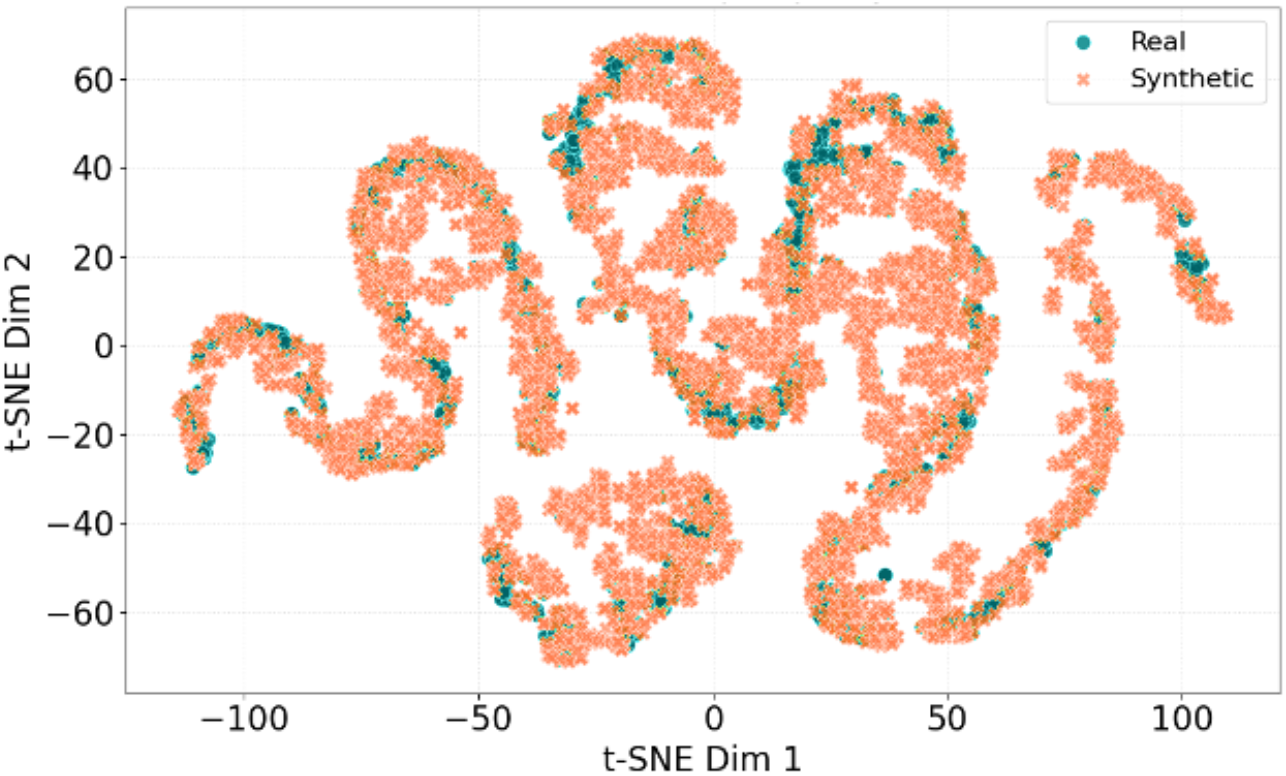
Two-dimensional t-SNE projection of real and synthetic metabolic trajectories. This visualization maps the high-dimensional Energy Expenditure (EE) of 5000 synthetic sequences onto a 2D manifold to evaluate structural fidelity at the trajectory level.

### Temporal fidelity of synthetic EE trajectories

While the earlier analyses showed the population-level statistical fidelity consistency of the EICD framework, a critical validation aspect for any time-series generative model is its ability to reproduce the temporal dynamics of individual physiological sequences. Figure 3 presents three randomly selected 30-minute synthetic EE trajectories generated by the full EICD model (ℳ_*full*_), each conditioned on a distinct metabolic context, and plotted together with their equivalent real empirical trajectory and the corresponding dynamic physiology guardrail corridor. These representative examples showed that across the three trajectories, the synthetic EE remained strictly within the physiology guardrail corridor, confirming a BVR of 0% at the individual trajectory level.

**Figure 3.**
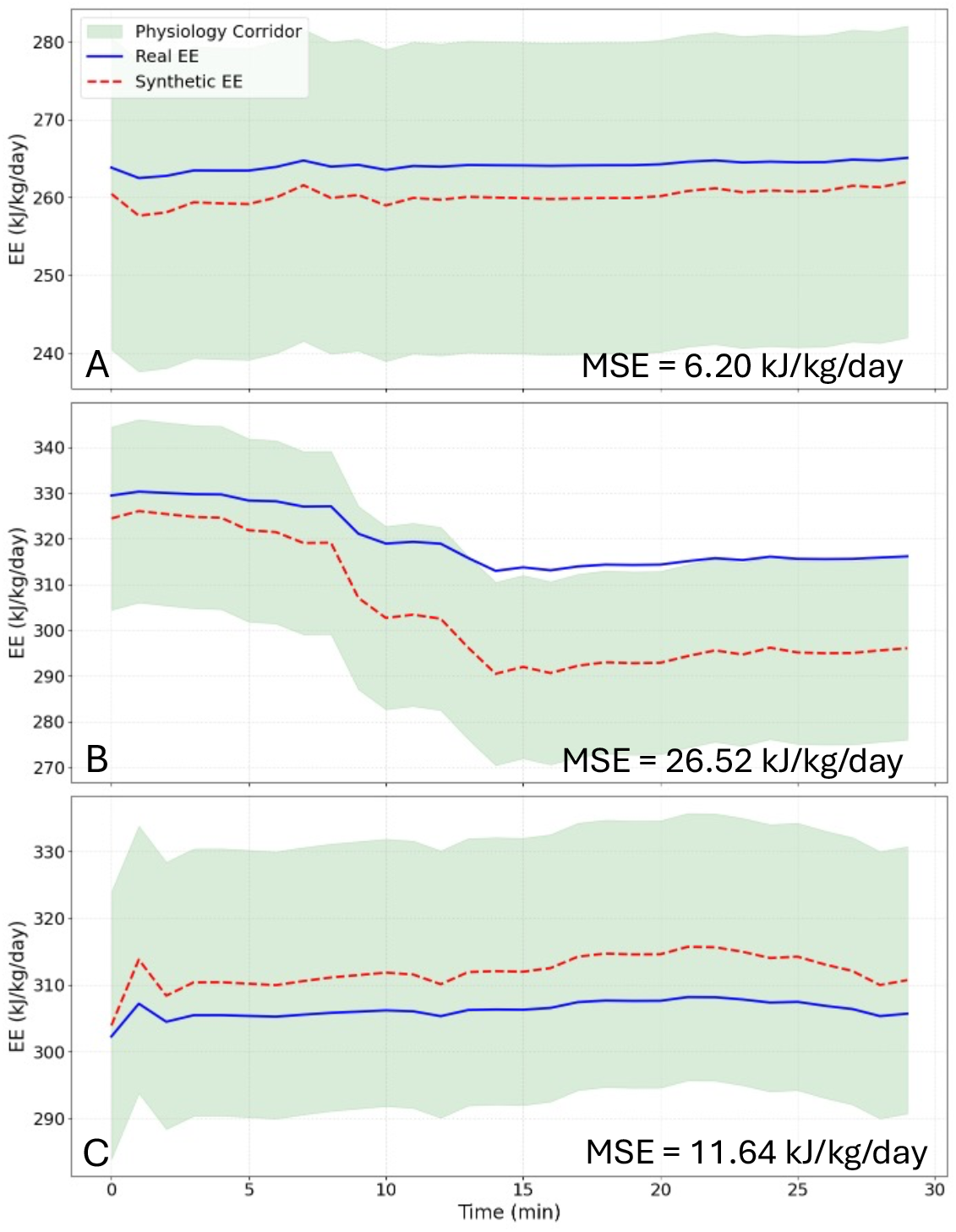
Three (A, B, and C) representative 30-minute real (blue solid lines) versus EICD-synthetic (red dashed lines) energy expenditure (EE) trajectories across their corresponding physiological guardrail corridor (shaded area).

While the MSE measures point-to-point deviation it ignores temporal structure, therefore, the temporal fidelity of the generated trajectories was quantified using the DTW distance the normalised cross-correlation. Averaged throughout the 5000 synthetic trajectories, the EICD framework achieved a mean MSE of 29 kJ/kg/day, a mean DTW distance of 23 kJ/kg/day, a peak cross-correlation coefficient of 0.84, and a mean temporal lag of +0.91 minutes. These results indicate that the EICD framework generates synthetic EE trajectories that are not only close in magnitude to the real signal but also structurally correct in shape and temporally well-aligned, with the synthetic response leading the real signal by less than one minute on average. The DTW distance of 23 kJ/kg/day, lower than the MSE of 29 kJ/kg/day, reveals that a portion of the apparent magnitude error captured by MSE is attributable to minor temporal misalignment rather than true metabolic divergence. When the time axis is optimally warped, the two signals align considerably more closely, confirming that the EICD framework captures the correct temporal dynamics of porcine energy expenditure even when instantaneous point-to-point relatively is imperfect. The cross-correlation coefficient of 0.84 further supports this interpretation, indicating high temporal synchrony between real and synthetic trajectories through different metabolic conditions. The mean positive lag of +0.91 minutes suggests that the synthetic signal marginally anticipates the real metabolic response, a behaviour consistent with the GRU-based denoising backbone learning a smoothed representation of the conditioning inputs that attenuates short-duration transient fluctuations while preserving the dominant temporal trend.

### Evaluation of physiological manifold consistency

Beyond statistical accuracy, the main advantages of the EICD framework lies in its adherence to physiological and bioenergetic laws. This subsection evaluates the physiological consistency of the generated trajectories (time series), specifically analysing how well the model respects the boundaries defined by the physiology guardrail.

As shown in Table 4, the ℳ_*full*_ model achieved a BVR of 0.93%, indicating that nearly all generated trajectories remain within the expert-defined metabolic limits of the physiology guardrail. This resulted BVR is closely approaches the 0.62% violation rate observed on the real data, which represents the noise floor of the empirical dataset.

On the other hand, the ℳ_*no_phys*_ variant resulted in a BVR of 12.10%. This large violation rate reveals a reliability problem, indicating that without the ℒ_*phys*_ constraint, the diffusion model frequently hallucinates metabolic states that are physically/physiologically impossible, such as *EE* falling below the allometric basal floor.

The *t*-SNE manifold overlap in Figure 2 further supports the physiological integrity of the EICD framework. Because the synthetic trajectories cover the exact same spatial manifold as the real observations, it can be concluded that the model is not just generating random valid numbers, but is capturing the complex, non-linear dependencies between physical activity (*ODBA*), environmental drivers (*T*_*a*_), and metabolic output. This confirms that the EICD framework produces a synthetic population that is not only statistically indistinguishable from the real one but is also mechanistically grounded.

Unlike standard generative models that may treat variables independently, the EICD framework must respects the dynamic relationship where energy expenditure is simultaneously governed by both the thermal environment (*T*_*a*_) and physical movement (*ODBA*). To quantify the framework’s ability to generate data that respects these complex interrelations of animal metabolism, we evaluated the alignment of the 3D metabolic manifolds (Figure 4A). As demonstrated in the overlapping 3D manifolds (Figure 4A), the synthetic model successfully reconstructed the non-linear metabolic topography observed in the empirical study. The localized performance of the model was further investigated through a physiological residual (Δ*EE* = *EE*_*Real*_ - *EE*_*Synth*_) map (Figure 4B). This spatial analysis reveals that predictive error is uniformly distributed across the primary metabolic manifold. The majority of the *T*_*a*_ and *ODBA* feature space exhibits residuals within ±3 kJ/kg/day, represented by the neutral white and light-coloured regions within Figure 3B. Quantitatively, the synthetic manifold achieved a 3D surface mean absolute error (MAE) of 9.33 kJ/kg/day relative to empirical benchmarks (Table 5). While minor diverging residuals (red/blue extremes) are visible at the manifold’s boundaries, specifically at the intersection of extreme cold (*T*_*a*_ < 12°C) and peak activity, these regions correspond to areas where individual biological variations are naturally most pronounced. Importantly, these boundary regions also represent the lowest-density areas of the empirical data distribution, meaning their proportional impact on overall model fidelity is limited. This is further verified by the *t*-SNE manifold coverage score of 87%, where the uncovered 13% is spatially consistent with the boundary residual regions indicated earlier. These results indicate that inherent individual variability represents the primary generative constraint at the manifold’s edge, despite the application of the physiology guardrail. Collectively, these metrics provide a mutually coherent validation of the model’s performance, identifying precisely where the model performs well and where the remaining generative limitations lies.

**Figure 4.**
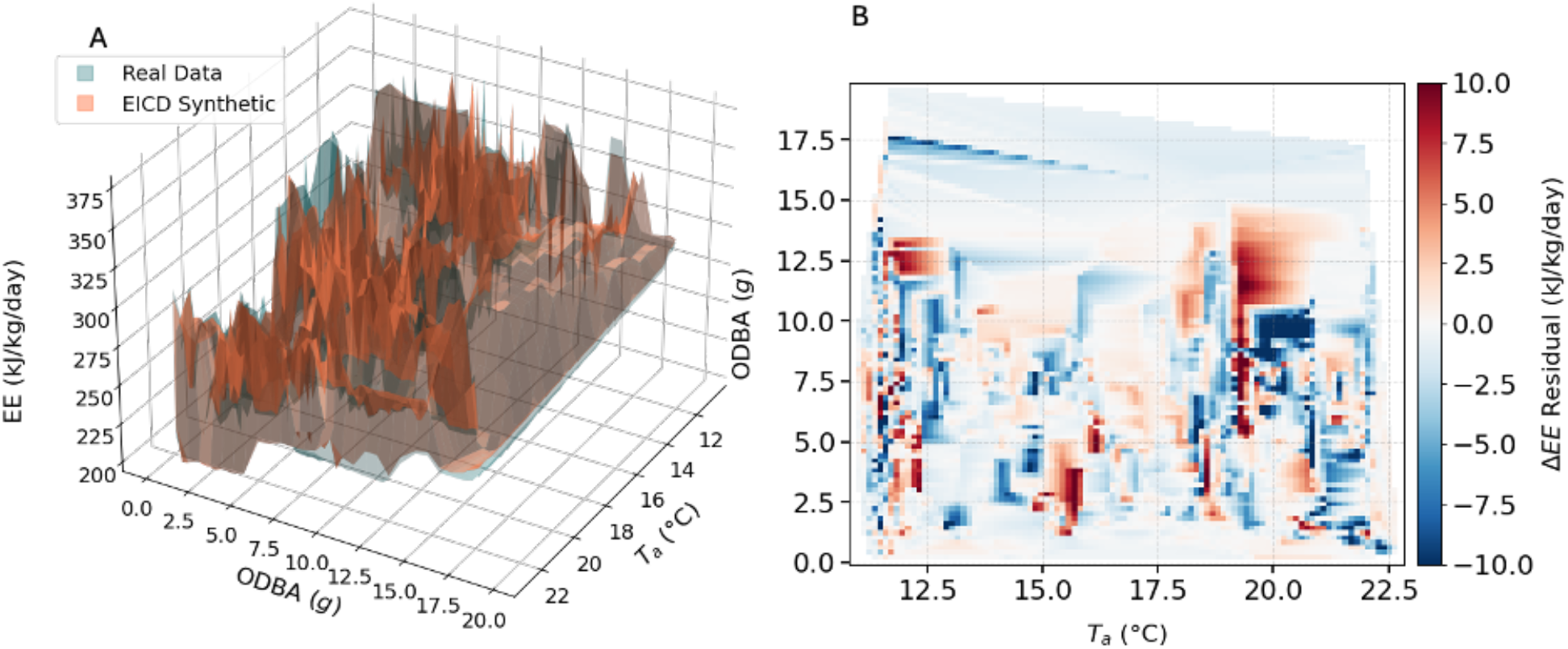
Comparative 3D manifold analysis of metabolic states. (A) three-dimensional metabolic manifold demonstrating the overlapping surface topology of empirical observations (Real Data) and EICD-generated sequences (Synthetic). The surfaces map energy expenditure (EE) as a function of ambient temperature (T_a_) and physical activity (ODBA). (B) Corresponding 2D residual heatmap (ΔEE = EE_Real_ - EE_Synth_) projecting the predictive accuracy across the operational environmental and behavioral envelope.

**Table 5.**
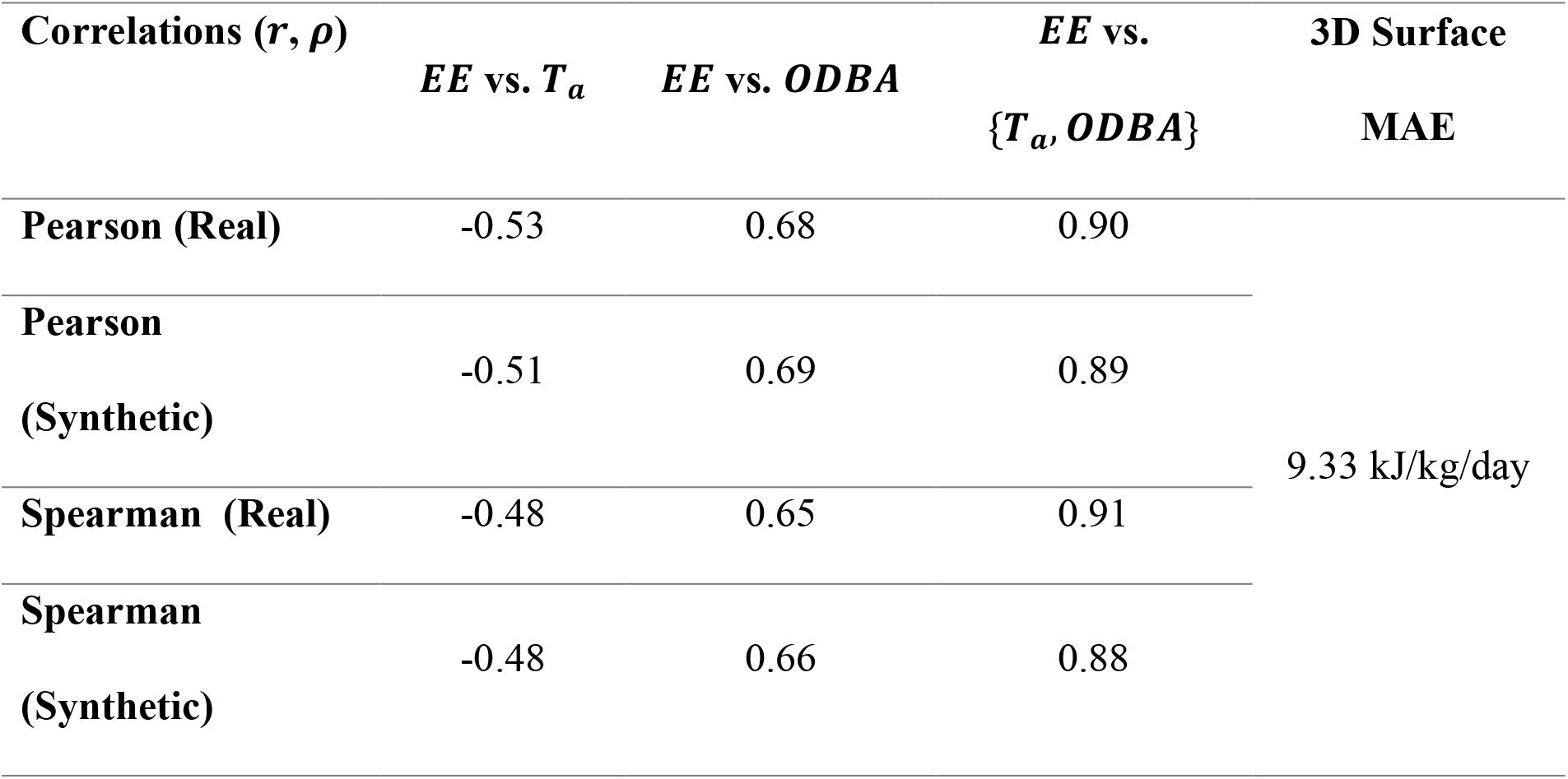
Comparative correlation matrix and global error analysis.

Furthermore, the EICD framework was validated by comparing the Pearson correlation coefficients of the primary metabolic drivers, *T*_*a*_ and *ODBA*, between the real and synthetic datasets (Table 5). The model successfully preserved the negative correlation between *EE* and *T*_*a*_ (real: -0.53; synthetic: -0.51) and the positive correlation between *EE* and activity levels was maintained (real: 0.68; synthetic: 0.69). Critically, the joint influence of temperature and activity on energy expenditure showed near-perfect alignment (real: 0.90; synthetic: 0.89). It is acknowledged that Pearson (*r*) is a linear association metric and may not fully capture the non-linear dependencies inherent in the metabolic manifold. To complement this, Spearman rank correlation coefficients (*ρ*) are reported in Table 5 as a non-linear dependence measure.

### The ablation study: insights and interpretations

The ablation study isolates the specific impact of the physiology guardrail, derived by ℒ_*phys*_, and the conditioning matrix (**c**) on the model’s generative fidelity. By systematically removing these components, we demonstrate that biological consistency is not an emergent property of standard diffusion models but requires explicit mechanistic constraints.

- *Impact of the physiology guardrail*: The removal of the physiological loss term, in the ℳ_*no_phys*_ variant, resulted in a significant degradation of biological integrity, with the BVR increasing from 0.93% to 12.10% (Table 4). This 13-fold increase in violations occurred despite the model maintaining relatively low statistical divergence (JSD = 0.090), confirming that statistical fidelity alone is insufficient to prevent hallucinated metabolic states.
- *Significance of dynamic conditioning*: The model variant ℳ_*uncond*_, which lacks the primary metabolic drivers resulted the most severe performance collapse, with a BVR spike to 29.43% and a KLD of 2.13 (Table 4). This highlights that porcine energy expenditure is a forced physiological response to ambient temperature and activity rather than an autonomous stochastic pattern.
- *Statistical fidelity, physiological integrity trade-off*: The full EICD framework variant (ℳ_*full*_) model achieved the optimal balance, providing high statistical fidelity (MSE = 29) while reaching a violation rate (0.93%) that aligns with the inherent noise floor of real-world empirical data (0.62%) as shown in Table 4.

### Implications for animal digital twin development

These results demonstrate that for animal digital twin applications, statistical accuracy is not a substitute for physiological viability. Hence, the development of the EICD framework marks a significant advancement toward high-fidelity animal digital twins (ADTs). By integrating mechanistic physiological knowledge with deep generative modeling, this research addresses several critical bottlenecks in the digitization of animal experimentation and animal digital twin development.

- *Enhancing predictive in silico expe*riments: The EICD framework ensures that synthetic agents (digital twins) operate within a physiologically consistent manifold. This capability allows researchers to conduct what-if simulations (Moingeon et al., 2023), such as predicting metabolic responses to extreme climate conditions or evaluating the effect of different feeding regimes, with high degree of confidence that the model will not generate biologically erroneous outliers.
- *Data augmentation and ethical animal research*: In many animal studies, data collection is limited by the cost of high-precision sensors and ethical constraints regarding animal handling (Lovarelli et al., 2023; Youssef et al., 2024). The ability to generate physiology-consistent synthetic trajectories provides a scalable solution for data augmentation especially longitudinal and time-series data. These synthetic datasets can be used to train downstream machine learning models (e.g., early disease detection or heat stress alarms) without the need for a massive influx of new empirical data. By refining the accuracy of synthetic models, we reduce the total number of animals required for invasive metabolic studies. This aligns with the global trend towards more adherence with the 3Rs in animal research.
- *Continuous monitoring and decision support*: By capturing how an animal’s energy expenditure changes in response to its surroundings, the EICD framework provides a robust engine for real-time decision support system (Badreldin et al., 2024). Digital twins can be synchronized with live sensor streams to monitor individual animal health and welfare continuously. For instance, when real-world physiological response diverges from the EICD-modeled normal manifold, the system can trigger automated alerts for immediate intervention. This capability transforms animal management from reactive to a predictive intervention, ensuring that deviations in health or welfare status are identified before they escalate.

### Challenges and limitations

While the EICD framework demonstrates significant fidelity in replicating metabolic signatures, its implementation highlights fundamental constraints in ADT development. These limitations primarily arise from the inherent trade-offs between generative flexibility, the model’s ability to produce diverse novel data, and biological precision, which requires strict adherence to rigid physiological laws.

- *Individual variations vs. regularization*: As shown in the physiological residual map (Figure 3B), predictive errors are disproportionately concentrated at the intersection of extreme environmental stressors and peak physical exertion (e.g., maximal shivering thermogenesis during acute cold exposure combined with high locomotor activity). While the global 3D surface MAE of 9.33 kJ/kg/day (Table 5) is low, these localized edge cases represent a significant challenge for simulating rare, high-intensity metabolic events. In biological systems, which are inherently non-linear and characterized by individual variability, responses at these boundary conditions become highly stochastic and varied (Gómez-Prado et al., 2022; Youssef, 2014). Consequently, the model faces a fundamental trade-off, it must replicate the natural volatility and individual-specific responses of biological data without violating the mechanistic guardrails imposed by species-specific metabolic laws.
- *Physiological adaptation and temporal drift*: Furthermore, a critical limitation of the current framework is its static nature regarding physiological adaptation. In real-world environments, animals undergo acclimatization, a process where prolonged exposure to a specific thermal or nutritional state leads to a shift in the basal metabolic rate and lower thermoneutral zone (Gómez-Prado et al., 2022; Wijffels et al., 2024). Because the EICD framework is currently trained on discrete physiological snapshots, it may lack the temporal depth required to simulate these gradual, long-term shifts in energy efficiency. This lack of “adaptive memory” could result in temporal drift, where the digital twin’s predictions remain accurate for acute responses but diverge from the animal’s actual state as it matures or acclimatizes to seasonal environmental changes (Schauberger et al., 2019).
- *Energy balance completeness*: The upper bound *U*(*c*) in the current framework accounts for basal metabolism, activity-induced thermogenesis, and thermoregulatory costs, but does not explicitly model anabolic energy costs (e.g., muscle growth, tissue deposition) or reproductive and production energy partitioning (e.g., milk synthesis, egg production) that are highly relevant in commercial animal production systems. Additionally, interaction terms between concurrent stressors (e.g., simultaneous heat stress and high activity) are not explicitly captured. This simplification is deliberate and appropriate for the growing-pig proof-of-concept context presented here. The EICD framework is, however, designed to be extensible, where field experts are encouraged to augment *U*(*c*) with additional energy terms relevant to their specific application and species, underscoring the philosophy that the expert defines the physiological boundaries that are most appropriate for their context.
- *Conditioning dependency*: The high BVR (29.43%) observed in the unconditional model variant proves that biological plausibility is strictly tied to the accuracy of input conditioning (*T*_*a*_ and *ODBA*). This dependency implies that the framework’s integrity is dependent on the precision of the used sensors. Furthermore, it is important to note that in commercial agricultural applications, the bioenergetic parameters used in this framework (e.g., *k*_*basal*_, thermoneutral zone limits, activity coefficients) are not static but inherently time-varying, shifting with growth stage, season, health status, and diet. The current implementation uses fixed expert-defined values, which is an intentional simplification for this proof-of-concept study focused on growing pigs under controlled conditions. The framework is explicitly designed so that field experts can substitute their own application-specific, time-varying parameter sets as warranted by their target context. Future work should address the integration of adaptive parameter schedules to improve long-term fidelity in dynamic production environments. Regarding temporal scope: the current framework generates 30-minute trajectories, a window deliberately chosen because it is sufficient to capture discrete physiological events of practical importance in precision livestock farming, such as a complete heat stress episode, a feeding bout, or an acute activity surge. For longer-horizon generation, a sliding-window chaining approach can be applied, where successive 30-minute windows are generated sequentially using the last conditioning state of the preceding window as input; however, it is acknowledged that small cumulative errors in predicted energy balance can lead to metabolic drift over extended periods, where the digital twin’s state slowly diverges from the animal’s actual physiological trajectory as the animal grows and acclimatizes. Addressing this long-term drift remains an important direction for future work.

## Conclusions and Future Work

This article presented the development and validation of the Expert-Informed Conditional Diffusion (EICD) framework, a novel generative approach to synthesizing high-fidelity metabolic time-series trajectories for animal digital twins (ADTs). Motivated by the critical data scarcity challenge in commercial animal production systems, where the continuous, high-resolution physiological monitoring required to build robust ADTs remains both economically prohibitive and ethically constrained, the EICD framework addresses the fundamental limitation of conventional generative models that prioritize statistical pattern-matching over biological reality, frequently producing physiological hallucinations that render synthetic data unsuitable for downstream livestock management applications.

The EICD framework demonstrates that the integration of expert-informed constraints, specifically through the physiology loss function (PhLF), is essential for capturing the complex, non-linear dynamics of porcine bioenergetics. The model achieved near-perfect distributional fidelity, characterized by an average Jensen-Shannon divergence (JSD) of 0.062 and a Kullback-Leibler divergence (KLD) of 0.19. Quantitative evaluation showed that the full EICD model achieved a mean energy expenditure (EE) of 284.94 ± 38.70 kJ/kg/day, mirroring the empirical average of 281.33 ± 41.58 kJ/kg/day with negligible deviation.

The results revealed that the physiology guardrail acts as a filter for generative quality, while standard diffusion models optimize solely for the likelihood of the data distribution, EICD penalizes samples that contradict expert-defined constraints, such as the porcine bioenergetics. This is quantitatively evidenced by the biological violation rate (BVR), where the full EICD model (ℳ_*full*_) significantly outperformed the unconditional variant (ℳ_*uncond*_), which exhibited a high BVR of 29.43%. These findings indicate that the physiological guardrail does not restrict generative diversity but instead steers the model toward a realistic physiological manifold, effectively preventing the distributional drift seen in models like ℳ_*no_phys*_. By providing a reliable method for generating physiology-consistent synthetic data conditioned on real production variables (ambient temperature, physical activity, and body weight), this framework directly addresses the data scarcity bottleneck that currently limits the development of robust animal digital twins in commercial livestock systems. The approach supports the 3Rs principles in animal research while simultaneously advancing the computational infrastructure needed for next-generation precision livestock farming, enabling extensive *in silico* experimentation that reduces the reliance on invasive metabolic trials.

### Future research directions

While this work establishes the efficacy of the EICD framework as a proof-of-concept for growing pigs under controlled thermal conditions, several opportunities for future investigation remain critical for translating this approach into deployable precision livestock farming tools. A primary direction is the extension of the framework to broader commercial production contexts, including broiler and laying hen systems, dairy cattle, and aquaculture species, each of which involves distinct bioenergetic laws, housing configurations, and production stressors that the expert-informed conditioning matrix must be adapted to reflect. Within the porcine domain, the current framework should be extended to encompass the full production cycle from weaning to slaughter weight, incorporating the dynamic growth-related shifts in metabolic baseline and thermoneutral zone that characterise commercial fattening operations. Biological systems are not static and exhibit significant acclimatization over time, necessitating future research to incorporate adaptive memory into the diffusion process to simulate how an animal’s metabolic baseline shifts during growth cycles. Expanding the conditioning manifold to include secondary production stressors such as relative humidity, air velocity, stocking density, and social competition will increase the model’s reliability in commercial environments where multiple stressors operate simultaneously and interact non-linearly.

From a diagnostic standpoint, a major objective for future work is the integration of causal reasoning to transform the EICD framework from a generative data augmentation tool into a real-time diagnostic engine capable of distinguishing between, for instance, thermal discomfort and sub-clinical disease onset, a distinction of direct practical relevance to farm managers seeking to minimise antibiotic use and reduce production losses. To enhance the robustness of the expert-defined physiological constraints across diverse species and production systems, future work will explore the use of retrieval-augmented generation (RAG) to autonomously extract and integrate species-specific bioenergetic laws and breed-specific metabolic parameters directly from technical literature, breed standards, and nutritional requirement databases. This would substantially reduce the expert parameterisation burden and facilitate rapid deployment across new production contexts. Finally, the computational efficiency of the denoising process must be optimised for real-time deployment on edge-computing devices within commercial animal houses, enabling instantaneous welfare alerts triggered by divergence between the live sensor stream and the digital twin’s predicted physiological manifold, a capability that would represent a significant step toward fully autonomous, AI-driven livestock welfare monitoring at farm scale.

## References

Aarts, Y., Vodorezova, K., Tang, Y., Heetkamp, M., Laurenssen, B., & Youssef, A. (2024). The EnergyTag: A Wearable Software Sensor for Online Monitoring of Animal’s Dynamic Energy Expenditure. In D. Berckmans, P. Tassinari, & D. Torreggiani (Eds.), 11th European Conference on Precision Livestock Farming (pp. 1307–1315). EA-PLF.

Badreldin, N., Cheng, X., & Youssef, A. (2024). An Overview of Software Sensor Applications in Biosystem Monitoring and Control. Sensors 2024, Vol. 24, Page 6738, 24(20), 6738. 10.3390/S24206738

Barré-Sinoussi, F., & Montagutelli, X. (2015). Animal models are essential to biological research: issues and perspectives. Future Sci. OA, 1(4), FSO63. 10.4155/fso.15.63

Bartali, E., Jongebreur, A., & Moffitt, D. (1999). CIGR handbook of agricultural engineering, Volume 2: Animal production and aquacultural engineering.

Berckmans, D. (2013). Basic principles of PLF: Gold standard, labelling and field data. Precision Livestock Farming 2013, 21–55. https://kuleuven.limo.libis.be/discovery/fulldisplay?docid=lirias1633686&context=SearchWebhook&vid=32KUL_KUL:Lirias&lang=en&search_scope=lirias_profile&adaptor=SearchWebhook&tab=LIRIAS&query=any,contains,LIRIAS1633686&offset=0

Brophy, E., Wang, Z., She, Q., & Ward, T. (2023). Generative Adversarial Networks in Time Series: A Systematic Literature Review. ACM Computing Surveys, 55(10). 10.1145/3559540

Butler, P. J., Green, J. A., Boyd, I. L., & Speakman, J. R. (2004). Measuring metabolic rate in the field: The pros and cons of the doubly labelled water and heart rate methods. Functional Ecology, 18(2), 168–183. 10.1111/J.0269-8463.2004.00821.X;PAGEGROUP:STRING:PUBLICATION

Chen, X., Roberts, R., Liu, Z., & Tong, W. (2023). A generative adversarial network model alternative to animal studies for clinical pathology assessment. Nature Communications 2023 14:1, 14(1), 7141-. 10.1038/s41467-023-42933-9

Friston, K. (2009). The free-energy principle: a rough guide to the brain? Trends in Cognitive Sciences, 13(7), 293–301. 10.1016/j.tics.2009.04.005

Gelfand, A. E., & Smith, A. F. M. (1990). Sampling-Based Approaches to Calculating Marginal Densities. Journal of the American Statistical Association, 85(410), 398. 10.2307/2289776

Ghosheh, G. O., Li, J., & Zhu, T. (2024). A Survey of Generative Adversarial Networks for Synthesizing Structured Electronic Health Records. ACM Computing Surveys, 56(6). 10.1145/3636424

Gómez-Prado, J., Pereira, A. M. F., Wang, D., Villanueva-García, D., Domínguez-Oliva, A., Mora-Medina, P., Hernández-Avalos, I., Martínez-Burnes, J., Casas-Alvarado, A., Olmos-Hernández, A., Ramírez-Necoechea, R., Verduzco-Mendoza, A., Hernández, A., Torres, F., & Mota-Rojas, D. (2022). Thermoregulation mechanisms and perspectives for validating thermal windows in pigs with hypothermia and hyperthermia: An overview. Frontiers in Veterinary Science, 9, 1023294. 10.3389/fvets.2022.1023294

Goodfellow, I., Pouget-Abadie, J., Mirza, M., Xu, B., Warde-Farley, D., Ozair, S., Courville, A., & Bengio, Y. (2020). Generative adversarial networks. Communications of the ACM, 63(11), 139–144. 10.1145/3422622

Halsey, L. G. (2016). Terrestrial movement energetics: current knowledge and its application to the optimising animal. The Journal of Experimental Biology, 219(Pt 10), 1424–1431. 10.1242/jeb.133256

Heetkamp, M. J. W., Alferink, S.. J. J., Zandstra, T., Hendriks, P., Brand, H. van den, & Gerrits, W. J. J. (2015). Design of climate respiration chambers, adjustable to the metabolic mass of subjects in: Indirect calorimetry. In In Indirect Calorimetry: Techniques, computations and applications (pp. 35–56). Wageningen Academic Publishers.

Higham, C., Higham, D. J., & Grindrod, P. (2025). Diffusion Models for Generative Artificial Intelligence: An Introduction for Applied Mathematicians. Society for Industrial and Applied Mathematics, 67(3), 607–623. 10.1137/23M1626232

Ho, J., Jain, A., & Abbeel, P. (2020). Denoising diffusion probabilistic models. In H. Larochelle, M. Ranzato, & R. T. Hadsell (Eds.), Proceedings of the 34th International Conference on Neural Information Processing Systems (pp. 6840–6851). Curran Associates Inc. https://dl.acm.org/doi/abs/10.5555/3495724.3496298

Karniadakis, G. E., Kevrekidis, I. G., Lu, L., Perdikaris, P., Wang, S., & Yang, L. (2021). Physics-informed machine learning. Nature Reviews Physics 2021 3:6, 3(6), 422–440. 10.1038/s42254-021-00314-5

Keogh, E., & Ratanamahatana, C. A. (2005). Exact indexing of dynamic time warping. Knowledge and Information Systems, 7(3), 358–386. 10.1007/S10115-004-0154-9/METRICS

Leiva, B., & Schramski, J. R. (2022). On the rules of life and Kleiber’s law: the macroscopic relationship between materials and energy. Heliyon, 8(6), e09647. 10.1016/j.heliyon.2022.e09647

Listou Ellefsen, A., Bjørlykhaug, E., Æsøy, V., Ushakov, S., & Zhang, H. (2023). Controlled physics-informed data generation for deep learning-based remaining useful life prediction under unseen operation conditions. Mechanical Systems and Signal Processing, 197, 110359. 10.1016/j.ress.2018.11.027

Loni, M., Poursalim, F., Asadi, M., & Gharehbaghi, A. (2025). A review on generative AI models for synthetic medical text, time series, and longitudinal data. Npj Digital Medicine 2025 8:1, 8(1), 281-. 10.1038/s41746-024-01409-w

Lovarelli, D., Bacenetti, J., & Guarino, M. (2023). Digital Twins in agriculture: challenges and opportunities for environmental sustainability. Current Opinion in Environmental Sustainability, 61, 101252. 10.1016/j.jclepro.2020.121409

Lu, M., Norton, T., Youssef, A., Radojkovic, N., Fernández, A. P., & Berckmans, D. (2018). Extracting body surface dimensions from top-view images of pigs. International Journal of Agricultural and Biological Engineering, 11(5), 182–191. 10.25165/IJABE.V11I5.4054

Maaten, L. van der, & Hinton, G. (2008). Visualizing Data using t-SNE. Journal of Machine Learning Research, 9(86), 2579–2605. http://jmlr.org/papers/v9/vandermaaten08a.html

Moingeon, P., Chenel, M., Rousseau, C., Voisin, E., & Guedj, M. (2023). Virtual patients, digital twins and causal disease models: Paving the ground for in silico clinical trials. Drug Discovery Today, 28(7), 103605. 10.1016/J.DRUDIS.2023.103605

National Research Council (NRC). (2012). Nutrient Requirements of Swine: Eleventh Revised Edition. In Nutrient Requirements of Swine (11th ed.). National Academies Press. 10.17226/13298

Panagiotakopoulos, T., Kotsiantis, S., Gkillas, A., & Lalos, A. S. (2025). Conditional Diffusion Models: A Survey of Techniques, Applications, and Challenges. IEEE Access, 13, 183617– 183643. 10.1109/ACCESS.2025.3625094

Peña Fernández, A., Demmers, T. G. M., Tong, Q., Youssef, A., Norton, T., Vranken, E., & Berckmans, D. (2019). Real-time modelling of indoor particulate matter concentration in poultry houses using broiler activity and ventilation rate. Biosystems Engineering, 187, 214–225.

Raissi, M., Perdikaris, P., & Karniadakis, G. E. (2019). Physics-informed neural networks: A deep learning framework for solving forward and inverse problems involving nonlinear partial differential equations. Journal of Computational Physics, 378, 686–707. 10.1016/j.jcp.2018.10.045

Schauberger, G., Mikovits, C., Zollitsch, W., Hörtenhuber, S. J., Baumgartner, J., Niebuhr, K., Piringer, M., Knauder, W., Anders, I., Andre, K., Hennig-Pauka, I., & Schönhart, M. (2019). Global warming impact on confined livestock in buildings: efficacy of adaptation measures to reduce heat stress for growing-fattening pigs. Climatic Change 2019 156:4, 156(4), 567–587. 10.1007/s10584-019-02525-3

Scholander, P. F., Hock, R., Walters, V., Johnson, F., & Irving, L. (1950). Heat regulation in some arctic and tropical mammals and birds. The Biological Bulletin, 99(2), 237–258. 10.2307/1538741

Schrama, J. W., Verstegen, M. W. A., Verboeket, P. H. J., Schutte, J. B., & Haaksma, J. (1996). Energy metabolism in relation to physical activity in growing pigs as affected by type of dietary carbohydrate. Journal of Animal Science, 74(9), 2220–2225. 10.2527/1996.7492220x

Speakman, J. R. (1999). The Cost of Living: Field Metabolic Rates of Small Mammals. Advances in Ecological Research, 30(C), 177–297. 10.1016/S0065-2504(08)60019-7

Tannenbaum, J., & Bennett, B. T. (2015). Russell and Burch’s 3Rs Then and Now: The Need for Clarity in Definition and Purpose. Journal of the American Association for Laboratory Animal Science: JAALAS, 54(2), 120. /pmc/articles/PMC4382615/

van Veen, L. A., van den Brand, H., van den Oever, A. C. M., Kemp, B., & Youssef, A. (2025). An adaptive expert-in-the-loop algorithm for flock-specific anomaly detection in laying hen production. Computers and Electronics in Agriculture, 229(6), 109755. 10.1016/j.compag.2024.109755

Wang, J., Upadhyay, D., Zaman, M., & Srikantha, P. (2025). Synthetic Power Flow Data Generation Using Physics-Informed Denoising Diffusion Probabilistic Models. 2025 IEEE International Conference on Communications, Control, and Computing Technologies for Smart Grids, SmartGridComm 2025 - Proceedings. 10.1109/SmartGridComm65349.2025.11204605

Wijffels, G., Lees, A. M., Sullivan, M. L., Sammes, S. L., Li, Y., & Gaughan, J. B. (2024). Allostasis as a consequence of high heat load in grain-fed feedlot cattle. Translational Animal Science, 8. 10.1093/tas/txae133

Yi, H., Hou, L., Jin, Y., Saeed, N. A., Kandil, A., & Duan, H. (2024). Time series diffusion method: A denoising diffusion probabilistic model for vibration signal generation. Mechanical Systems and Signal Processing, 216, 111481. 10.1016/j.ymssp.2024.111481

Youssef, A. (2014). Model-Based Control of Micro-Environment with Real-Time Feedback of Bioresponses. KU Leuven.

Youssef, A. (2023). Soft Sensor and Biosensing. Encyclopedia of Smart Agriculture Technologies, 1–10. 10.1007/978-3-030-89123-7_171-1

Youssef, A., Jansen, C., & Neethirajan, S. R. (2022). Soft-Sensing Approach for Predicting Bovine Respiratory Disease Severity. Precision Livestock Farming ‘22, 932–939. https://research.wur.nl/en/publications/soft-sensing-approach-for-predicting-bovine-respiratory-disease-s

Youssef, A., Norton, T., & Berckmans, D. (2019). Bioenvironmental Zonal Controlling of Incubated Avian Embryo Using Localised Infrared Heating. Processes, 7(10), 651.

Youssef, A., Peña Fernández, A., Wassermann, L., Biernot, S., Wittauer, E.-M., Bleich, A., Hartung, J., Berckmans, D., & Norton, T. (2020). An Approach towards Motion-Tolerant PPG-Based Algorithm for Real-Time Heart Rate Monitoring of Moving Pigs. Sensors, 20(15), 4251.

Youssef, A., Viazzi, S., Exadaktylos, V., & Berckmans, D. (2013). Control system for positioning of broilers using infrared heating and thermal camera. Precision Livestock Farming 2013 - Papers Presented at the 6th European Conference on Precision Livestock Farming, ECPLF 2013.

Youssef, A., Vodorezova, K., Aarts, Y., Agbeti, W. E. K., Palstra, A. P., Foekema, E., Aguilar, L., Torres, R. da S., & Grübel, J. (2024). IUMENTA: A generic framework for animal digital twins within the Open Digital Twin Platform. http://arxiv.org/abs/2411.10466

Zheng, H., Yang, Z., Liu, W., Liang, J., & Li, Y. (2015). Improving deep neural networks using softplus units. Proceedings of the International Joint Conference on Neural Networks, 2015-September. 10.1109/IJCNN.2015.7280459

Zhou, J., Li, J., Zheng, G., Wang, X., & Zhou, C. (2024). MTSCI: A Conditional Diffusion Model for Multivariate Time Series Consistent Imputation. International Conference on Information and Knowledge Management, Proceedings, 3474–3483. 10.1145/3627673.3679532

